# Human cytomegalovirus attenuates AKT activity by destabilizing insulin receptor substrate proteins

**DOI:** 10.1101/2023.04.17.537203

**Authors:** Anthony J. Domma, Felicia D. Goodrum, Nathaniel J. Moorman, Jeremy P. Kamil

## Abstract

The phosphoinositide 3-kinase (PI3K)/AKT pathway plays crucial roles in cell viability and protein synthesis and is frequently co-opted by viruses to support their replication. Although many viruses maintain high levels of AKT activity during infection, other viruses, such as vesicular stomatitis virus and human cytomegalovirus (HCMV), cause AKT to accumulate in an inactive state. To efficiently replicate, HCMV requires FoxO transcription factors to localize to the infected cell nucleus (Zhang et. al. mBio 2022), a process directly antagonized by AKT. Therefore, we sought to investigate how HCMV inactivates AKT to achieve this. Subcellular fractionation and live cell imaging studies indicated that AKT failed to recruit to membranes upon serum-stimulation of infected cells. However, UV-inactivated virions were unable to render AKT non-responsive to serum, indicating a requirement for *de novo* viral gene expression. Interestingly, we were able to identify that *UL38* (pUL38), a viral activator of mTORC1, is required to diminish AKT responsiveness to serum. mTORC1 contributes to insulin resistance by causing proteasomal degradation of insulin receptor substrate (IRS) proteins, such as IRS1, which are necessary for the recruitment of PI3K to growth factor receptors. In cells infected with a recombinant HCMV disrupted for *UL38*, AKT responsiveness to serum is retained and IRS1 is not degraded. Furthermore, ectopic expression of UL38 in uninfected cells induces IRS1 degradation, inactivating AKT. These effects of UL38 were reversed by the mTORC1 inhibitor, rapamycin. Collectively, our results demonstrate that HCMV relies upon a cell-intrinsic negative feedback loop to render AKT inactive during productive infection.

## IMPORTANCE

HCMV requires inactivation of AKT to efficiently replicate yet precisely how AKT is shut off during HCMV infection has remained unclear. We show that UL38, an HCMV protein that powerfully activates mTORC1, is necessary and sufficient to destabilize IRS1, a model insulin receptor substrate (IRS) protein. Degradation of IRS proteins in settings of excessive mTORC1 activity is a key mechanism for insulin resistance. When IRS proteins are destabilized, PI3K cannot be recruited to growth factor receptor complexes and hence AKT membrane recruitment, a rate limiting step in its activation, fails to occur. Despite its penchant for remodeling host cell signaling pathways, our results reveal that HCMV relies upon a cell-intrinsic negative regulatory feedback loop to inactivate AKT. Given that pharmacological inhibition of AKT potently induces HCMV reactivation from latency, our findings also imply that UL38 expression must be tightly regulated within latently infected cells to avoid spontaneous reactivation.

## INTRODUCTION

As obligate intracellular pathogens, viruses frequently remodel host-cell signal transduction pathways to suit their own interests, for instance, to evade innate and cell-mediated immunity and to harness macromolecular synthesis resources toward the production of infectious progeny virions. The phosphoinositide 3-kinase (PI3K)/ AKT (protein kinase B) pathway is targeted by a wide array of DNA viruses, such as Vaccinia virus, hepatitis B virus, herpes simplex virus-1 [1–3], as well as RNA viruses, e.g., vesicular stomatitis virus (VSV), measles virus, and human immunodeficiency virus [4–6]. However, in some examples AKT activity is maintained during lytic replication, while in other examples, such as VSV and HCMV, AKT accumulates in an inactive form in infected cells. Given that AKT plays crucial roles in cell viability, metabolism, and macromolecular synthesis [7,8], why certain viruses require its inactivation to efficiently replicate is unclear.

As summarized in **FIG 1A**, the PI3K/AKT pathway is canonically activated by cell surface receptors, e.g., G-protein coupled receptors and receptor tyrosine kinases (RTKs), in response to extracellular stimuli such as growth factors and chemokines. A key event during growth factor signaling is the recruitment of insulin receptor substrate (IRS) proteins, such as IRS1, to the cytoplasmic domains of the cognate receptor tyrosine kinases. Briefly, the RTKs phosphorylate IRS proteins at key tyrosine residues, which enables the binding and membrane recruitment of src homology 2 (SH2) domain-containing proteins such as PI3K to the activated receptor complex [9,10] [11]. Ultimately, this leads to the activation of PI3K, which converts phosphoinositol 4,5-bisphosphate (PIP2) to phosphoinositol 3,4,5– trisphosphate (PIP3) within the cytoplasmic facing, inner leaflet of cell membranes [12]. Accumulation of PIP3 provides docking sites for additional signaling proteins that contain pleckstrin homology (PH)-domains, such as AKT and the phosphoinositide-dependent kinase 1 (PDK1) [13,14]. The mechanistic target of rapamycin (mTOR) complex 2, mTORC2, then phosphorylates AKT at Ser473 [15,16], a site within a hydrophobic motif at the C-terminus of AKT. Phosphorylation at Ser473 enables docking to PDK1, which phosphorylates AKT at Thr308 on the T-loop of the catalytic protein kinase core [17,18]. Together, these two phosphorylation events result in fully activated AKT, which then disassociates from the membrane and phosphorylates a large number of substrates to mediate various effects on cell physiology [7]. For instance, AKT phosphorylates tuberous sclerosis complex 2 (TSC2), relieving its inhibition of a crucial mTOR complex, mTORC1 [19–21]. Once active, mTORC1 phosphorylates many different substrates, including S6 kinase (S6K) and 4E-BP1 [22,23] leading to increased protein synthesis, a key downstream effect observed upon treatment of cells with growth factors, such as insulin or EGF. Forkhead box class O (FoxO’s) transcription factors (TFs) are also important substrates of AKT, and in this case, AKT negatively regulates their nuclear localization [24]. Briefly, when AKT is inactive, FoxO TFs localize to the nucleus, where they regulate genes necessary for stress responses and cell differentiation [25–27]. In settings where extracellular signals favor cell cycle progression, AKT phosphorylates FoxO TFs, enabling them to associate with 14-3-3 proteins that mediate their export from the nucleus [28].

**Figure 1:**
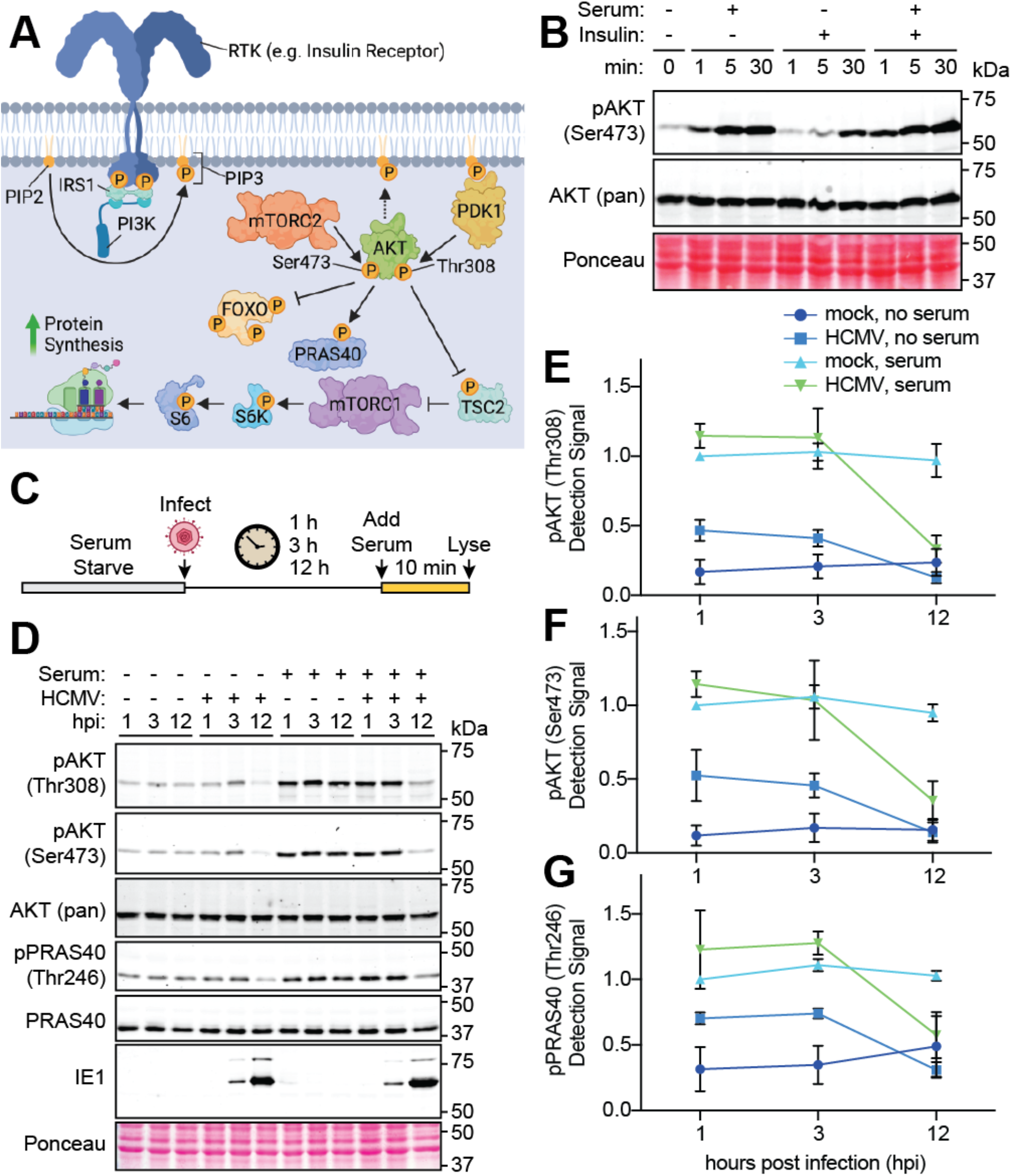
HCMV prevents AKT activation in response to serum. (**A**): Stimulation of growth factor receptor tyrosine kinases (RTK) causes the recruitment of the scaffolding adaptor insulin receptor substrate 1 (IRS1) to the receptor complex. IRS1 is then phosphorylated, which provides a docking site for recruitment of class IA phosphoinositol 3-kinases (PI3K). PI3K phosphorylates the lipid substrate phosphoinositol 4,5 diphosphate (PIP2) to generate phosphoinositol 3,4,5 triphosphate (PIP3). The pleckstrin homology (PH) domain of AKT binds to PIP3, which drives its localization to membranes where mTORC2 and phosphoinositol-dependent kinase 1 (PDK1) are available to phosphorylate AKT at Ser473 and Thr308, respectively, causing full activation its kinase activity. Once activated, AKT phosphorylates many substrates, such as the proline-rich AKT substrate of 40 kDa (PRAS40) at Thr246, Forkhead box class O (FoxO) family of transcription factors, and the tuberous sclerosis complex 2 protein (TSC2). Phosphorylation of TSC2 leads to activation of mTORC1, which in turn phosphorylates and activates S6 kinase (S6K). S6K phosphorylates S6 to increase protein synthesis. [Schematic generated in BioRender.] (**B**): Serum-starved fibroblasts were stimulated with insulin or 5% newborn calf serum for the indicated times (min) prior to lysis. Protein extracts were then resolved by SDS-PAGE, transferred to nitrocellulose membranes for Western blot analysis using a phospho-specific antibody to detect AKT phosphorylated at Ser473 compared to AKT (pan) antibody. (**C-G**): Fibroblasts were serum starved overnight and either mock infected or infected with HCMV strain Towne at multiplicity of infection (MOI=2 TCID50/cell) for the indicated times, h post-infection (hpi). Cells were then either treated (or mock-treated) with serum for 10 min prior to lysis for Western blot analysis. (**E-G**): Fluorescent signals from dye labeled secondary antibodies were quantified and phospho-specific antibody detection of pAKT Thr308, pAKT Ser473 and PRAS40 Thr246 was normalized to AKT (pan) or PRAS40, respectively. The arithmetic mean was calculated and graphed. Error bars indicate standard error of the mean (SEM) from three independent biological replicates. Where indicated, Ponceau staining was used visualize protein loading and confirm efficient transfer.

As with many cell signaling pathways, the PI3K/AKT circuit is subject to negative feedback, which has been most extensively studied in the context of insulin resistance. The literature has revealed a mechanism wherein mTORC1 phosphorylates IRS1 and IRS2 to cause their proteasomal degradation [29–34]. In brief, mTORC1 phosphorylation of IRS1 at Ser422 leads to its ubiquitination and proteasomal degradation [32]. When this crucial scaffolding adaptor is degraded, activation of PI3K ceases and AKT cannot be recruited to the membrane to be phosphorylated at Ser473 and Thr308, hence preventing its activation [35,36].

Human cytomegalovirus (HCMV), a betaherpesvirus that is an important cause of congenital defects in newborns and of life-threatening opportunistic infections in immunocompromised individuals [37–40], inactivates the PI3K/AKT pathway during productive infection [41–43]. However, little is known about which HCMV proteins interact with components of the PI3K/AKT pathway. The HCMV protein UL138 regulates EGFR signaling and is thought to promote AKT activation for the establishment of latency. Moreover, inhibition of PI3K/AKT signaling is observed to cause reactivation from latency in HCMV and several other herpesviruses [43,44]. UL7, a secreted protein, activates the PI3K/AKT pathway and subsequently inactivates FoxO family TFs in CD34+ progenitor cells, which is also thought to contribute to the establishment of latency [45]. Furthermore, the antiapoptotic protein UL38 reportedly activates mTORC1 both by direct activation [46,47] and by binding and inhibiting TSC2, a key negative regulator of mTORC1 [48]. Additionally, various virion envelope glycoprotein complexes bind to cell surface receptors to trigger activation of the PI3K/AKT pathway during viral entry into several different cell types [41,49–51]. We recently showed that HCMV infection causes AKT inactivation, which drives nuclear localization of FoxO TFs, and that artificially regulated nuclear localization of chimeric estrogen receptor FoxO3a fusion proteins rescue viral replication defects that otherwise occur during expression of constitutively-active AKT [42] which is in agreement with other reports [52]. These results indicated that HCMV relies upon the inactivation status of AKT to efficiently replicate. Nonetheless, how HCMV inactivates AKT has remained unclear.

The literature suggests that mTORC1 phosphorylation of IRS1 leads to its degradation in settings of excess growth factor signaling, resulting in dampened PI3K/AKT activity, a key mechanism that contributes to insulin resistance. Because the HCMV protein UL38 has been identified to be a particularly powerful activator of mTORC1, we investigated whether UL38 is required during infection to inactivate AKT. Indeed, our results demonstrate that UL38 is both necessary and sufficient to destabilize IRS1 and inactivate AKT. Conversely, we observe that pharmacological inhibition of mTORC1 during HCMV infection prevents IRS1 degradation, causing AKT to remain active during infection.

## RESULTS

### HCMV infection renders AKT unresponsive to serum

To investigate how HCMV modifies the responsiveness of AKT to growth factors, we first established the basic kinetics of AKT activation in both non-infected and HCMV infected and human fibroblasts, making use of phospho-specific antibodies to detect phosphorylation of AKT and of PRAS40, a canonical AKT substrate. Within 5 min of stimulating uninfected, (serum-starved) human fibroblasts with either serum-or insulin-stimulation, we detected increased phosphorylation of AKT at Ser473 (pAKT Ser473), a site relevant to AKT activation (**FIG 1B**). Moreover, this phosphorylation was maintained for at least 30 min. Because serum treatment led to increased detection of pAKT Ser473 relative to insulin, we opted to use serum instead of insulin in our subsequent experiments (**FIG 1B**). As depicted in **FIG 1C**, we infected serum-starved cells with HCMV strain Towne at a multiplicity of infection (MOI) of 2 tissue culture infectious dose 50% (TCID_50_) units per cell (**FIG 1C–G**). Then, at either 1 h, 3 h, or 12 h post-infection (hpi), we stimulated the cells with 5% calf serum for 10 min (**FIG 1C**), before harvesting lysates for Western blot analysis. Levels of AKT phosphorylated at Thr308 or Ser473, two sites that both must be phosphorylated for maximal kinase activity [16,17,53], remained low in the absence of serum, as did levels of PRAS40 phosphorylated at Thr246, an AKT-specific phospho-acceptor site (**FIG 1C**). However, these three sites were readily phosphorylated upon serum treatment in both mock-infected cells, and in cells that had been infected with HCMV for up to 3 h. By 12 hpi, however, AKT and PRAS40 had become refractory serum stimulation, as evidenced by our detection of phosphorylation levels similar to those seen from non-stimulated serum-starved cells (**FIG 1D**). Data from three independent biological replicates were quantified and normalized to cognate detection signals for total AKT and PRAS40 (**FIGS 1E–G**). Levels of the 72 kDa HCMV immediate-early protein, IE1, indicated comparable levels of infection across samples and uniform protein loading. From these results, we concluded that HCMV infection renders AKT non-responsive to serum stimulation, in agreement with our previous findings [42,43].

### HCMV blocks recruitment of AKT to membranes

We next sought to investigate why serum stimulation fails to activate AKT during HCMV infection. We hypothesized that HCMV either (i) encodes a protein that directly binds AKT to prevent its phosphorylation by mTORC2 and/or PDK1, or (ii) indirectly achieves this by preventing activation of PI3K. To distinguish between these possibilities, we carried out membrane fractionation using differential ultracentrifugation, which allows for density-dependent separation of membranes from cytoplasmic fractions following the mechanical lysis of cells.

As expected, in serum-starved, non-infected conditions AKT is poorly co-fractionated with membranes. In contrast, upon serum stimulation, we observed a 2-fold increase in levels of AKT associated with membranes of non-infected cells. (Of note, the depletion of the cytosolic enzyme GAPDH in membrane fractions indicated that our fractionation procedure was effective.) In HCMV-infected samples, however, only low levels of AKT were recovered from membrane fractions even during serum stimulation, resembling what was seen in membranes from serum-starved, non-infected cells (**FIG 2A–B**). From these results, we concluded that HCMV-infected cells show defects in AKT membrane recruitment in response to serum stimulation.

**Figure 2:**
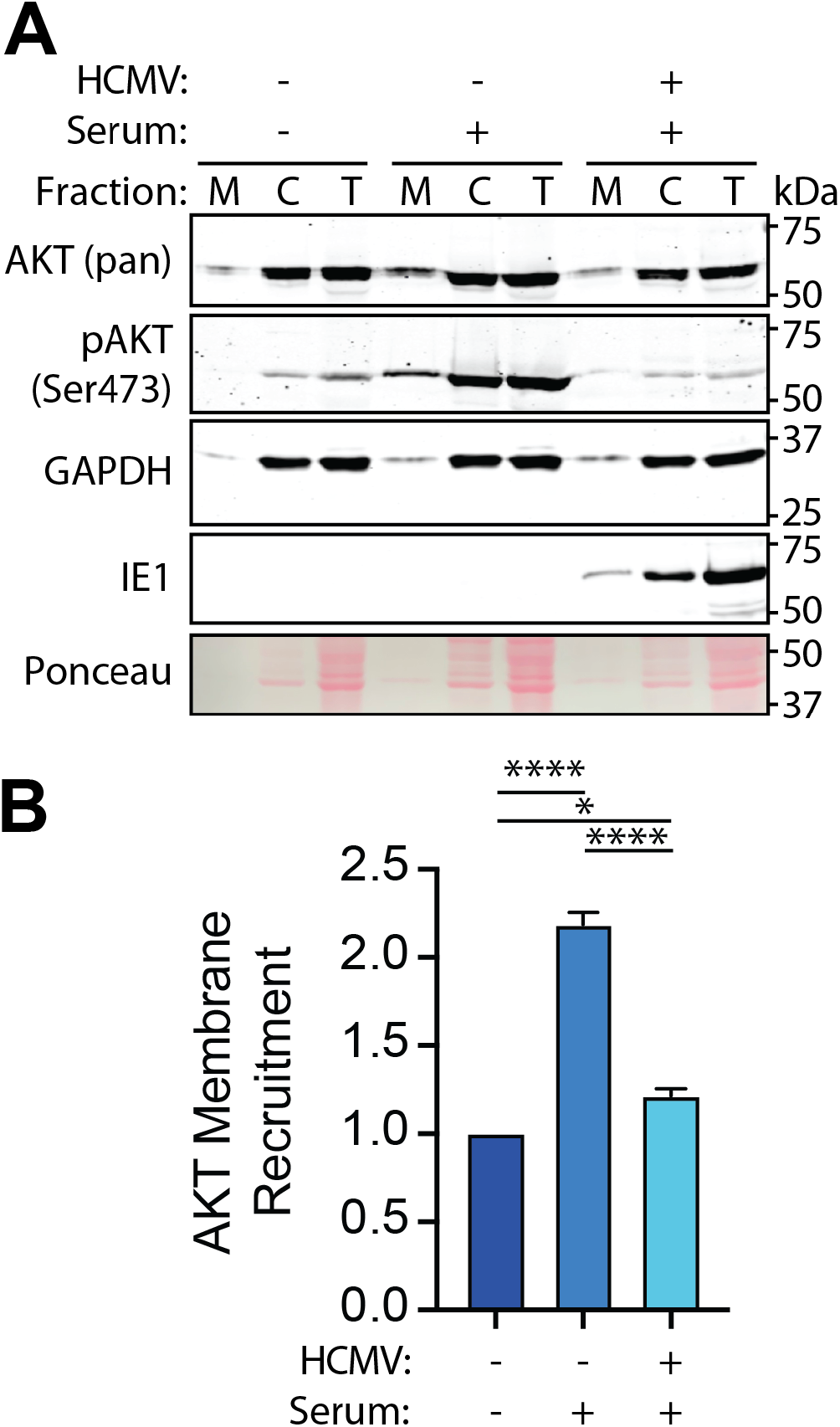
Membrane recruitment of AKT is defective in HCMV-infected cells. (**A-B**) Serum-starved fibroblasts were either mock-infected or infected with HCMV strain Towne (MOI=2 TCID50/cell) for 12 hours and either mock-treated or treated with 5% NCS (serum) for 10 minutes. Samples were mechanically lysed and differentially centrifuged to separate the membrane (M) fraction from the cytoplasmic (C) fraction and subjugated to Western blot analysis of indicated proteins alongside total (T) lysate. Ponceau staining was used as a loading control. (**B**) The AKT (pan) bands were quantified and the membrane band was normalized to the total lysate band. Quantification was then normalized to-Serum, the arithmetic mean was calculated and values were graphed. Error bars=SEM. n=4.

To further investigate AKT membrane recruitment, we conducted live-cell imaging studies of HeLa cells expressing the AKT pleckstrin homology (PH) domain fused to *A. victoria-*enhanced green fluorescent protein (GFP), as illustrated in **FIG 3A**. Because our results up to this point evaluated HCMV inactivation of AKT in fibroblasts, we first confirmed that HCMV also inactivates AKT during infection of serum-fed HeLa cells, an epithelial cell line that is permissive for HCMV infection, but which does not support efficient production of viral progeny [54]. We used strain AD169 repaired for *UL131* (AD169_r131) in these studies since the pentameric glycoprotein complex (gH/gL/gO/UL128-UL131) is needed for entry into epithelial cells [55,56]. Having observed that HCMV inactivates AKT within 12 h of infection in HeLa cells (similar to what we observed in fibroblasts in **FIG 5C**), we transfected HeLa cells with a plasmid expressing the PH domain of AKT genetically fused to GFP (AKT-PH(WT)-GFP), [where, WT: wild-type], or as a negative control, an R25C missense mutant that is defective for PIP3 binding (**FIG 3A**) [57]. The next day cells were serum-starved for 2 h prior to visualization by live cell imaging. In the presence of serum, we detected a 2-fold increase in GFP signal at the membranes of uninfected HeLa cells, as compared to serum-starved controls, matching the size of the effect we saw in our membrane fractionation studies of uninfected fibroblasts. In contrast, HCMV-infected HeLa cells showed poor membrane recruitment, exhibiting levels of GFP signal at the membrane that were similar to negative controls, despite treatment with serum (**FIG 2D–E**). Taken together with our membrane fractionation results (**FIG 2**), these findings further suggested defects in AKT membrane recruitment during HCMV infection.

**Figure 3:**
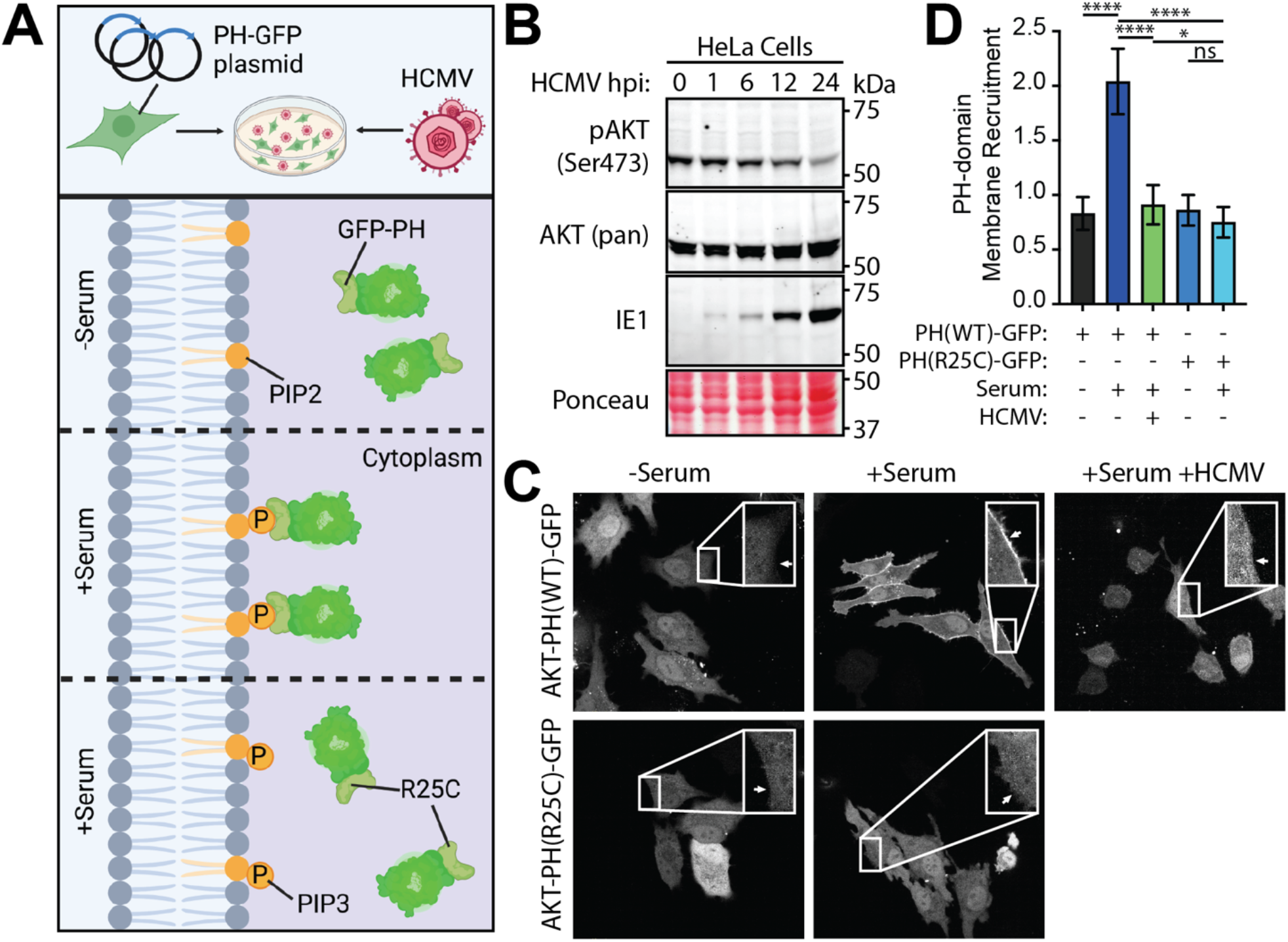
PH-domain fails to recruit to the membrane of HCMV-infected cells. (A) HeLa cells were transfected with a plasmid expressing GFP fused to the PH domain of AKT and infected with HCMV. In the absence of serum, PI3K signaling is shut off and no PIP3 is generated, so the PH domain of AKT remains in the cytoplasm. However, with the addition of serum, PI3K is activated and PIP3 is generated allowing PH domain binding to PIP3 and membrane recruitment. R25C mutants of the PH domain, even in the presence of serum, fail to bind to PIP3. Figure was created using BioRender. (B) HeLa cells were infected in the presence of 10% fetal bovine serum (FBS) with HCMV strain AD169r131 (MOI=2 TCID50/cell) and lysates were collected at the indicated hours post-infection (hpi). A Western blot analysis was performed probing for indicated proteins. Ponceau staining was used for loading control. n=1. (C-D) HeLa cells were transfected with an expression plasmid harboring the PH-domain of AKT fused to GFP or an R25C mutant of the AKT PH-domain fused to GFP as the transgene. Cells were then either infected or mock-infected with HCMV strain AD169r131 (MOI=2 TCID50/cell) for 12 hours. Cells were then serum-starved for 2 hours and either treated or mock treated with 10% FBS for 10 minutes. (D) Pixel density at the membrane was normalized to cytoplasmic pixel density and graphed. Tukey’s multiple comparisons statistical test was used to compare each condition to each other and labeled with asterisks (ns=P>0.05, *=P<0.05, **=P<0.01, ***=P<0.001, ****P<0.0001). n=3. Error bars=SEM.

**Figure 4:**
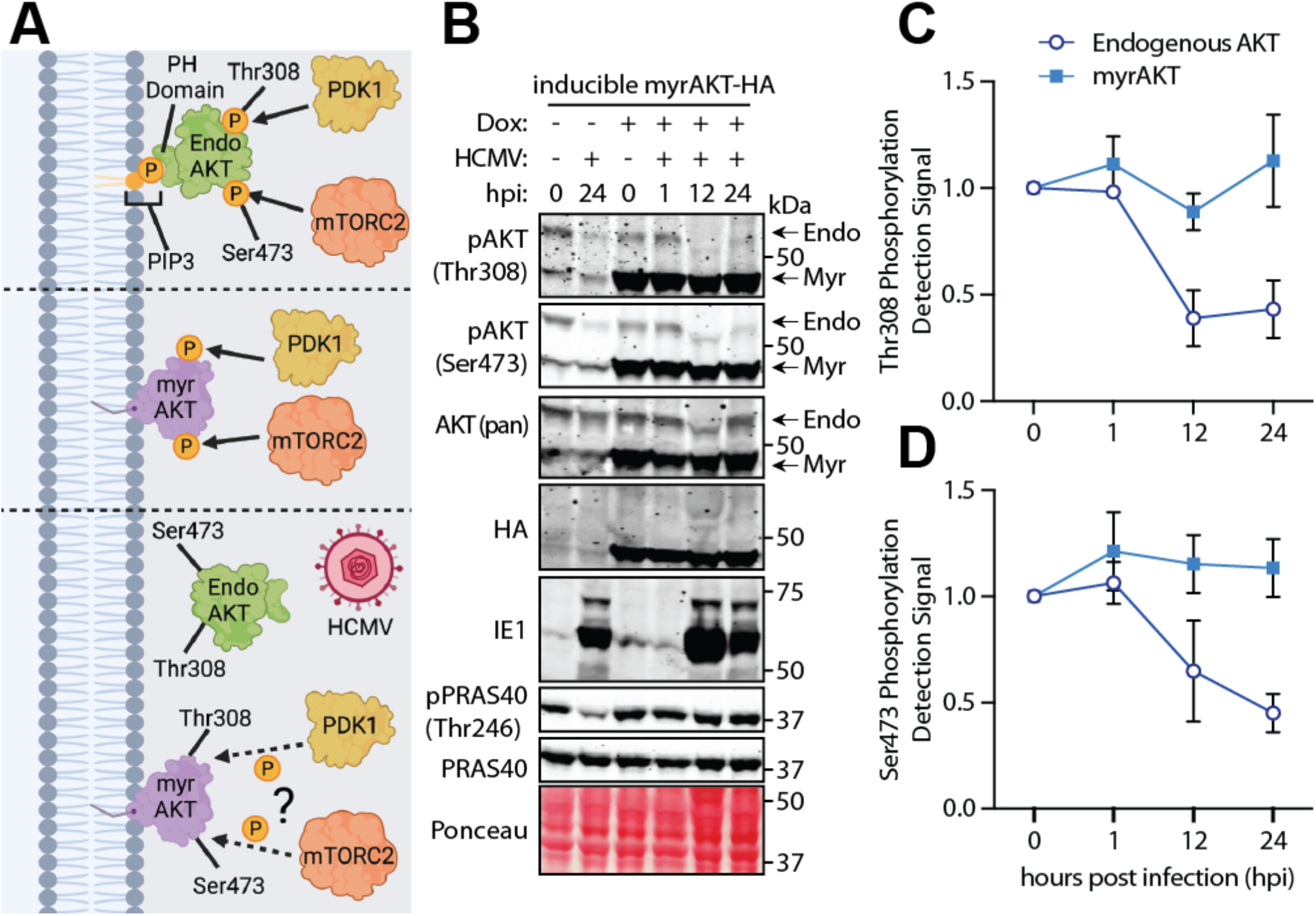
PDK1 and mTORC2 remain active during HCMV-mediated AKT inactivaiton. (**A**) PDK1 and mTORC2 phosphorylate AKT at Thr308 and Ser473 respectively following AKT membrane recruitment. Fibroblasts were transduced with a lentivirus harboring a “tet-on” promoter controlling expression of an AKT1 gene whose PH-domain has been replaced with a myristoylation (myr) signal and genetically fused to an hemagglutinin (HA) tag. Following expression, myrAKT is embedded in the membrane simulating constitutive “membrane recruitment” and is phosphorylated as such by PDK1 and mTORC2. This experiment addresses if the activators downstream of AKT, PDK1 and mTORC2, are active during HCMV infection. Figure was created using BioRender. (**B**) Induction of myrAKT occured after overnight treatment with doxycycline followed by infection or mock infection with HCMV strain TOWNE (MOI=2 TCID50/cell) for indicated hours post infection (hpi). A Western blot was performed probing for the indicated proteins using Ponceau stain as a readout for loading. In pAKT and Total AKT blots, both endogenously expressed (Endo) AKT and myrAKT (Myr) migrated to different molecular weights due to deletion of PH-domain in myrAKT gene. (**C**) pAKT (Thr308) signal and (**D**) pAKT (Ser473) signal were quantified, normalized to AKT (pan) and subsequently normalized to 0 hpi signal at respective molecular weight. The arithmetic mean was calculated and graphed. Error bars=SEM. n=3.

**Figure 5:**
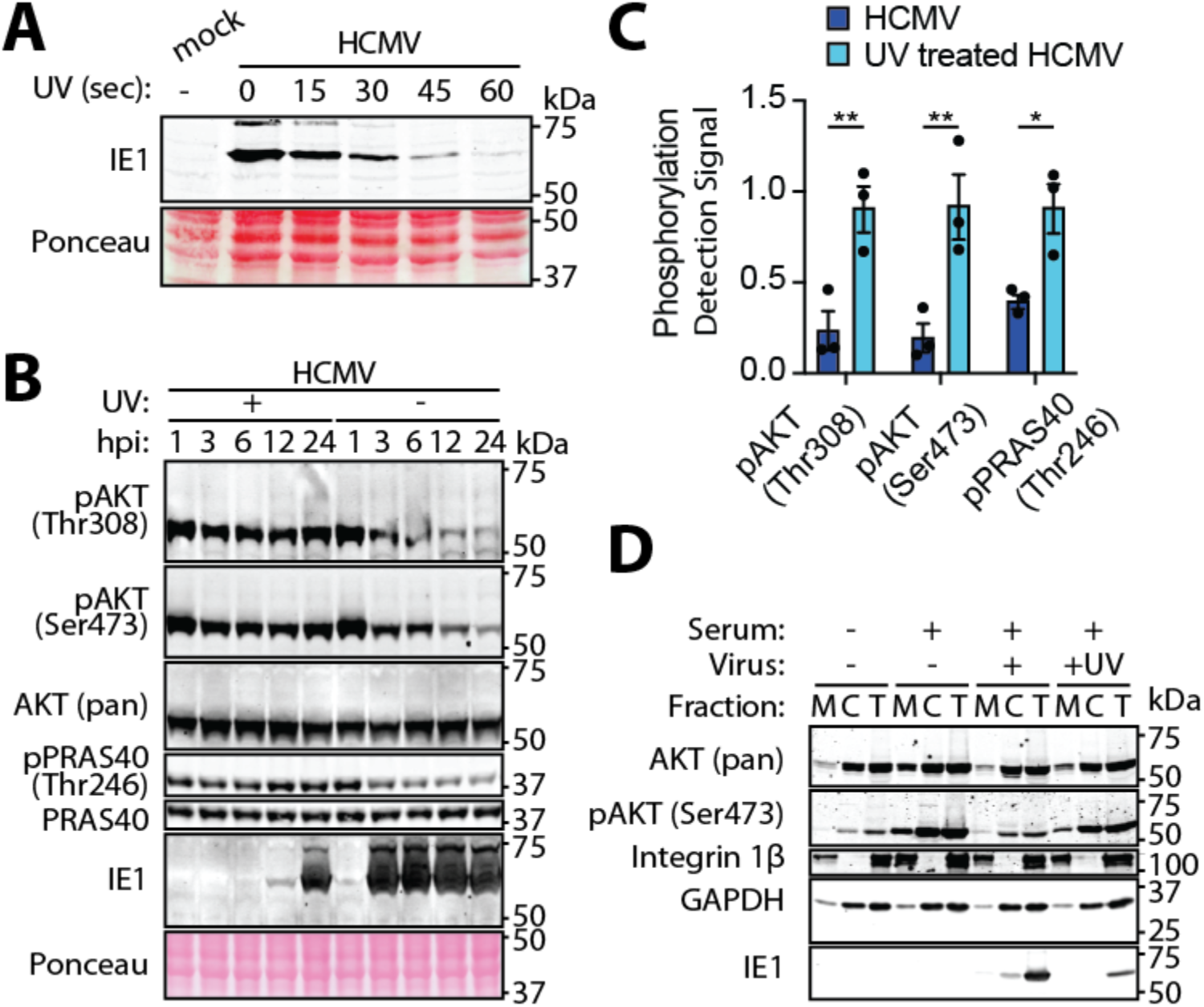
UV-treated virus fails to inactivate AKT. (**A**) HCMV strain Towne was mock-treated or treated with 125mJ of UV light for indicated seconds (sec) and allowed to infect fibroblasts (MOI=2 TCID50/cell) for 12 hours. A Western blot was performed staining for IE1 with Ponceau stain used as a readout for loading. n=1. (**B-C**) Fibroblasts were infected with mock-treated or UV-treated HCMV strain Towne (MOI=2 TCID50/cell) in the presence of serum and lysates were taken at the indicated hours post-infection (hpi). A Western blot was performed probing for the indicated proteins using Ponceau stain as a readout for loading. (**C**) Bands were quantified at 24 hpi, normalized to signal at 1 hpi, and the arithmetic mean was found and graphed. Sidak statistical test were used to compare each condition to the control and labeled with asterisks (ns=P>0.05, *=P<0.05, **=P<0.01, ***=P<0.001, ****P<0.0001). Error bars=SEM. n=3. (**D**) Serum-starved fibroblasts were either mock-infected or infected with UV-treated or untreated HCMV strain Towne (MOI=2 TCID50/cell) for 12 hours and either mock-treated or treated with 5% NCS (serum) for 10 minutes. Samples were mechanically lysed and differentially centrifuged to separate the membrane (M) fraction from the cytoplasmic (C) fraction and subjugated to Western blot analysis of indicated proteins alongside total (T) lysate.

### Loss of AKT signaling can be rescued by forcing its localization to membranes

We next sought to determine whether PDK1 and mTORC2, the key kinases that activate AKT subsequent to its localization to membranes [15,16,58], retained their ability to do so in HCMV-infected cells. To test this, we employed fibroblasts stably transduced with a “Tet-On” (doxycycline-inducible) myrAKT expression cassette, which encodes an AKT cDNA in which the PH domain is replaced with a retroviral myristoylation signal [42]. Upon doxycycline (Dox) treatment, myrAKT protein is expressed and targeted to membranes, regardless of whether PI3K is active (**FIG 4A**). After inducing the tet-on promoter for 24 h, cells were infected with HCMV in the presence of serum. A time course series of samples were harvested for Western blot analysis to assess the phosphorylation of AKT at Thr308 and Ser473 and of PRAS40 at Thr246. Because of their distinct relative mobilities in SDS-PAGE, endogenous AKT was readily distinguished from Dox-induced myrAKT, which was also fused to an HA-tag (**FIG 4B**). Similar to **FIG 6B** and previous publications, endogenous AKT from myrAKT-expressing cells was poorly reactive with Thr308 and Ser473 phospho-specific antibodies, despite the presence of serum [42,43]. However, the artificially expressed myrAKT remained phosphorylated at both Thr308 and Ser473 even during infection (**FIG 4B–D**). PRAS40 also remained phosphorylated at Thr246, a readout for AKT activity, in the myrAKT condition even during dephosphorylation at Ser473 and Thr308 of endogenous AKT. PRAS40 phosphorylation at Thr246 indicates that myrAKT remained active during HCMV-mediated AKT inactivation (**FIG 4B**). Identical experiments were performed, but using a K179M mutant, kinase-dead version of myrAKT, giving similar results (data not shown). Of note, in uninduced control conditions (where doxycycline was not added), weak detection of myrAKT protein gene product was observed, likely because the use of serum that had not been certified “free of tetracyclines” was used. Nonetheless, this small degree of leaky expression did not confound data interpretation because the regulation of endogenous AKT in HCMV-infected cells has been thoroughly characterized. Collectively, these results indicated to us that forcing AKT to localize to membranes rescued its phosphorylation and activation. Furthermore, these data also suggested that during HCMV infection AKT becomes refractory to activation downstream of serum stimulation due to a defect in upstream signaling that impairs AKT recruitment to membranes, not because PDK1 or mTORC2 activities are fundamentally lacking.

**Figure 6:**
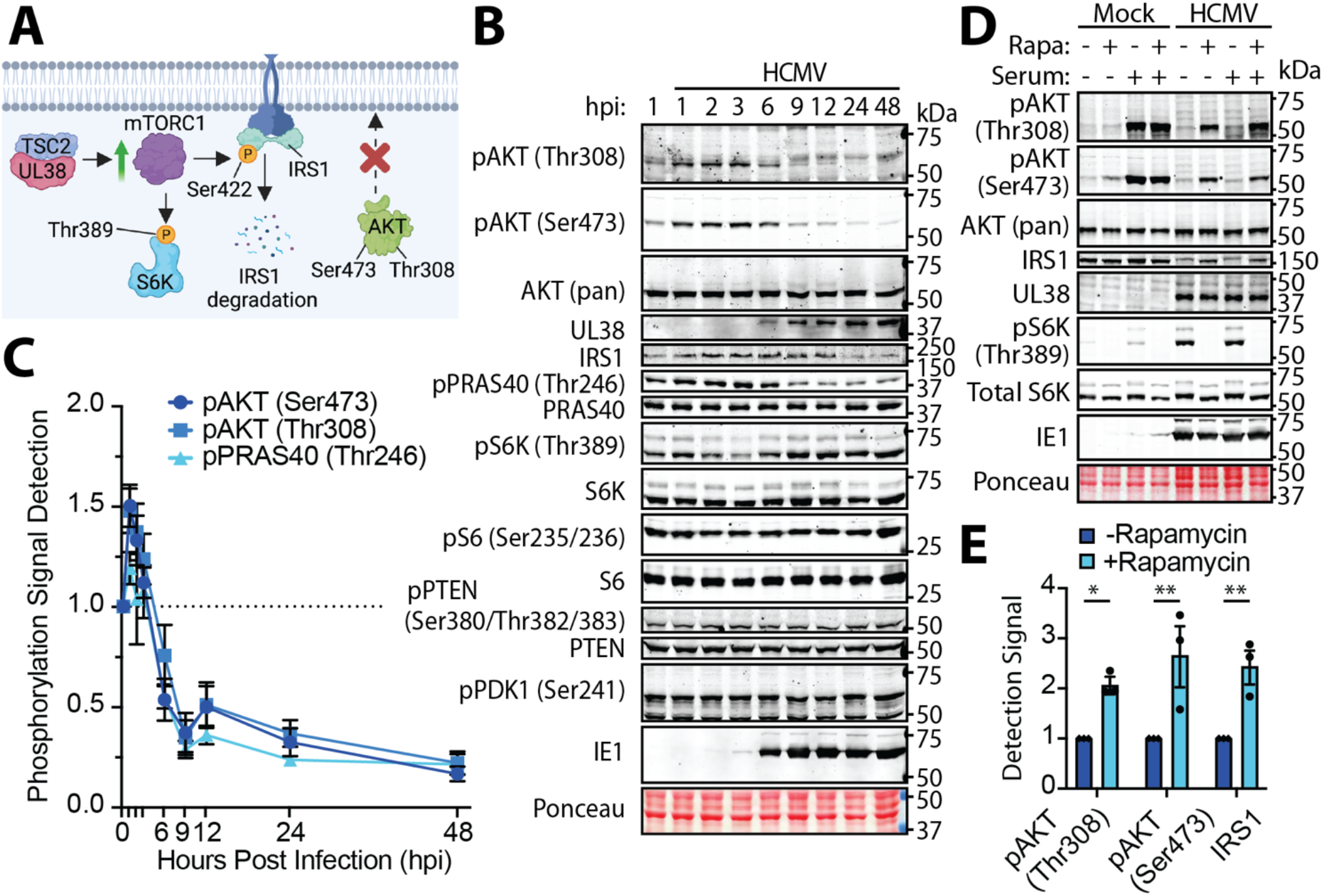
HCMV causes degradation of insulin receptor substrate 1. (**A**) Model for AKT inactivation in HCMV infected cells. HCMV UL38 mediated activation of mTORC1 leads to mTORC1 phosphorylation of IRS1, concomitantly inducing IRS1 degradation. This destabilization of IRS1 prevents PI3K recruitment to growth factor receptors (not shown), preventing AKT membrane recruitment and activation. Cartoon was generated using BioRender. (**B-C**) Fibroblasts were mock-infected or infected with HCMV strain Towne (MOI=2 TCID50/cell) in the presence of serum and lysates were taken at the indicated hours post-infection (hpi). A Western blot analysis detecting the indicated proteins was performed and (C) bands were quantified and phospho-specific quantification was normalized to total protein and arithmetic mean was found and graphed. The dotted line is the phosphorylation of mock-infected controls at 1 hpi, to which all infected time points are normalized to. Error bars=SEM. n=11. (**D-E**) Similar to Figure 1C, fibroblasts were serum starved overnight and either mock infected or infected with HCMV strain Towne multiplicity of infection (MOI=2 TCID50/cell). At 3 hpi, cells were mock-treated or treated with Rapamycin. At 24 hpi, cells were then either treated or mock-treated with serum for 10 minutes, and lysates were taken for Western blot analysis detection of indicated proteins. Ponceau staining was used for loading control. (**E**) Bands of the HCMV-infected cells were quantified, and phospho-specific quantification was normalized to total protein then normalized to ‘No Rapamycin’ control and arithmetic mean was found and graphed. Sidak statistical test were used to compare each condition to the control and labeled with asterisks (ns=P>0.05, *=P<0.05, **=P<0.01, ***=P<0.001, ****P<0.0001). Error bars=SEM. n=3.

### Viral gene expression is required to attenuate AKT recruitment to membranes and signaling

We next set out to identify which of the >185 genes present within the HCMV genome [59–61] encodes the factor responsible for AKT inactivation. To narrow the scope of genes implicated in AKT inactivation to either structural or non-structural genes, we first inactivated virions by treating virus preparations with ultraviolet (UV) light, which cross-links pyrimidine bases to form pyrimidine dimers in virion genomic DNA such that viral gene expression is severely impaired [62–64]. If AKT became refractory to serum stimulation after infection with a UV-inactivated virus, this would indicate that AKT inactivation does not require *de novo* viral gene expression, and hence would suggest that one or more virion-associated (“structural”) proteins are responsible. However, if AKT serum responsiveness was maintained after infection with UV-inactivated virions, this would indicate that *de novo* viral gene expression is required and that virion constituents, such as tegument proteins, are not sufficient to inactivate AKT.

After establishing that a 1 min exposure of virion preparations to 125 mJ of 254 nm radiation was sufficient to render IE1 protein expression virtually undetectable at 12 hpi (**FIG 5A**), we proceeded to infect fibroblasts, in the presence of serum, with either mock-treated or UV (125 mJ)-inactivated virions, collecting lysates for Western blot analysis to evaluate effects on phosphorylation of AKT and PRAS40. In the UV-inactivated virus condition, we observed that phosphorylation at key residues was rescued through 24 hpi for AKT (Thr308 and Ser473) and PRAS40, a canonical AKT substrate (Thr246) (**FIG 5B–C**). However, cells infected with untreated virus showed a greater loss of phosphorylation at all three phospho-acceptor sites, as expected (**FIG 6B)** [42,43]. As an orthogonal approach, we carried out membrane fractionation comparing fibroblasts infected with UV-inactivated virus versus untreated control virus. We observed increased co-fractionation of AKT integrin 1β, a transmembrane protein, that we used as a marker for faithful isolation of membrane fractions, in UV-inactivated virus-infected cells compared to untreated control infections (**FIG 5D**). Taken together, these results argue that *de novo* viral gene expression is required to render AKT defective for membrane recruitment and activation during infection.

### UL38 activation of mTOR induces degradation of IRS1

We next sought to identify the viral gene product that renders AKT refractory to serum stimulation and to gain further insights into the mechanism responsible for the defect in its membrane recruitment during infection. The literature argues for the existence of an important negative feedback loop within the PI3K/AKT/mTORC1 signaling pathway [8,32,65–67]. Briefly, aside from phosphorylating its downstream substrates, e.g., factors involved in mRNA translation, cell survival, and metabolism, mTORC1 also phosphorylates insulin receptor substrate (IRS) proteins, such as IRS1, to down modulate responsiveness to serum growth factors, a phenotype commonly referred to as “insulin resistance” [68]. mTORC1 phosphorylation of IRS family proteins (e.g., IRS1 at Ser422) results in their proteasomal degradation via the SCFβ-TRCP E3 ubiquitin ligase complex. Because IRS proteins play critical roles as adaptor proteins necessary for PI3K recruitment to the internal cytoplasmic domains of activated receptor tyrosine kinases, when they proteins are degraded AKT, in turn, also fails to recruit to membranes and thus cannot become activated in response to serum growth factors [32,35].

Notably, HCMV encodes a powerful activator of mTORC1, UL38 [46,48]. We, therefore, sought to test the hypothesis that UL38 is required for mTORC1-mediated phosphorylation and proteasomal degradation of IRS proteins during HCMV infection since this would be expected to render AKT refractory to serum stimulation (**FIG 6A**). Hence, we infected fibroblasts with HCMV strain Towne in the presence of serum and observed decreased IRS1 abundance and AKT phosphorylation over time by Western blots. By 24 hpi, we observed a 31% loss of IRS1 detection signal, accompanied by an IRS1 mobility shift similar to that observed by Yoneyama et al., which they attributed to mTORC1-mediated phosphorylation of IRS1 at Ser422 [32,68]. AKT phosphorylation initially showed a transient increase during infection, so detection of Ser473 and Thr308 phospho-epitopes peaked at 1 hpi. Nonetheless, by 9 hpi the detection signal for these phospho-sites declined to 25% and 23%, respectively, of their peak levels. Our results indicated that these AKT phospho-epitopes continued to decline over time. By 48 hpi, the final time point analyzed, the detection signal for Ser473 phosphorylation declined to 11% of peak levels, and Thr308 decreased to 15% relative to its 1 hpi peak. As expected, phosphorylation of PRAS40 (Thr246) decreased in tandem with that of AKT. Detection signals for phospho-PRAS (Thr246) decreased to 24% and 18% of its 1 hpi peak levels, at 9 hpi and 48 hpi, respectively (**FIG 6B–C**), consistent with our previous reports [42,43]. Notably, the kinetics with which AKT phosphorylation decreased during infection tightly correlated with the accumulation of UL38 protein (**FIG 6B**). Additionally, p70 S6K and ribosomal protein S6 remained phosphorylated at Thr389 [69] and Ser235/236 [70], respectively, which indicates that mTORC1 was active throughout the time series (**FIG 6B**). A previous report argued that PTEN, a dual protein and lipid phosphatase that serves as a key negative regulator of the PI3K/AKT pathway, is activated during HCMV infection, and the authors argued that this accounts for the loss of AKT activity in HCMV infected cells [71]. However, we were unable to reproduce this observation in studies with HCMV strain Towne-infected fibroblasts (**FIG 6B**), nor in endothelial or epithelial cells infected with pentamer-repaired strain AD169-infected (data not shown). Taken together, these results are consistent with the hypothesis that UL38-mediated activation of mTORC1 is required for proteasomal degradation of IRS1 during HCMV infection.

We next sought to establish whether inhibition of mTORC1 in HCMV-infected cells would prevent the degradation of IRS1 and restore AKT responsiveness to serum. To address this, we treated HCMV-infected cells with rapamycin, an mTORC1 kinase inhibitor, which would block the ability of mTORC1 to phosphorylate IRS1. Rapamycin is expected to prevent IRS1 degradation [68], and hence restore AKT serum responsiveness. In HCMV infected cells that had not been treated with rapamycin, we observed a magnificent increase in phosphorylation of S6K at Thr389, an mTORC1-mediated post-translational modification that is strongly stimulated by UL38 during infection [47,48]. As expected, however, rapamycin treatment abolished phosphorylation of S6K, such that pS6K (Thr 389) was undetectable in our hands. This confirmed that our rapamycin treatments were efficacious. At 24 hpi, we observed a 185% increase in IRS1 levels in protein extracts from rapamycin-treated infections compared to untreated infected controls. These effects were accompanied by increases in detection signals for AKT phospho-epitopes Thr308 (230%) and Ser473 (250%), relative to controls (**FIG 6D–E**). These results show that inhibition of mTORC1 rescues both IRS1 protein levels in HCMV-infected cells (just as classically observed in non-infected cell contexts) [68] and hence, restores serum responsiveness of AKT. Therefore, our findings suggest that mTORC1-mediated IRS1 degradation renders AKT refractory to serum stimulation during HCMV infection.

We next sought to address whether UL38 would be sufficient to cause IRS1 degradation and render AKT refractory to serum stimulation in non-infected cells. We thus carried out experiments in which serum-fed fibroblasts harboring doxycycline (Dox)-inducible expression cassette encoding *UL38* fused at its N-terminus to an influenza A virus hemagglutinin (HA) epitope tag, were Dox-induced to activate HA-UL38 expression. We then gathered lysates over a post-induction time course, evaluating for effects on IRS1 and AKT by Western blot. At 24 h post-Dox induction, we detected HA-tagged UL38 protein as well as pS6K (Ser389), confirming that the induced UL38 protein was biologically active (**FIG 7A**). Concomitantly, we observed a reduction in IRS1 levels, accompanied by a decrease in IRS1 mobility, similar to that seen in **FIG 6B** [32]. Additionally, a reduction in the detection signals for phospho-Thr308 and phospho-Ser473 AKT was evident in cells induced for HA-UL38 expression (**FIG 7A**), as expected [47].

**Figure 7:**
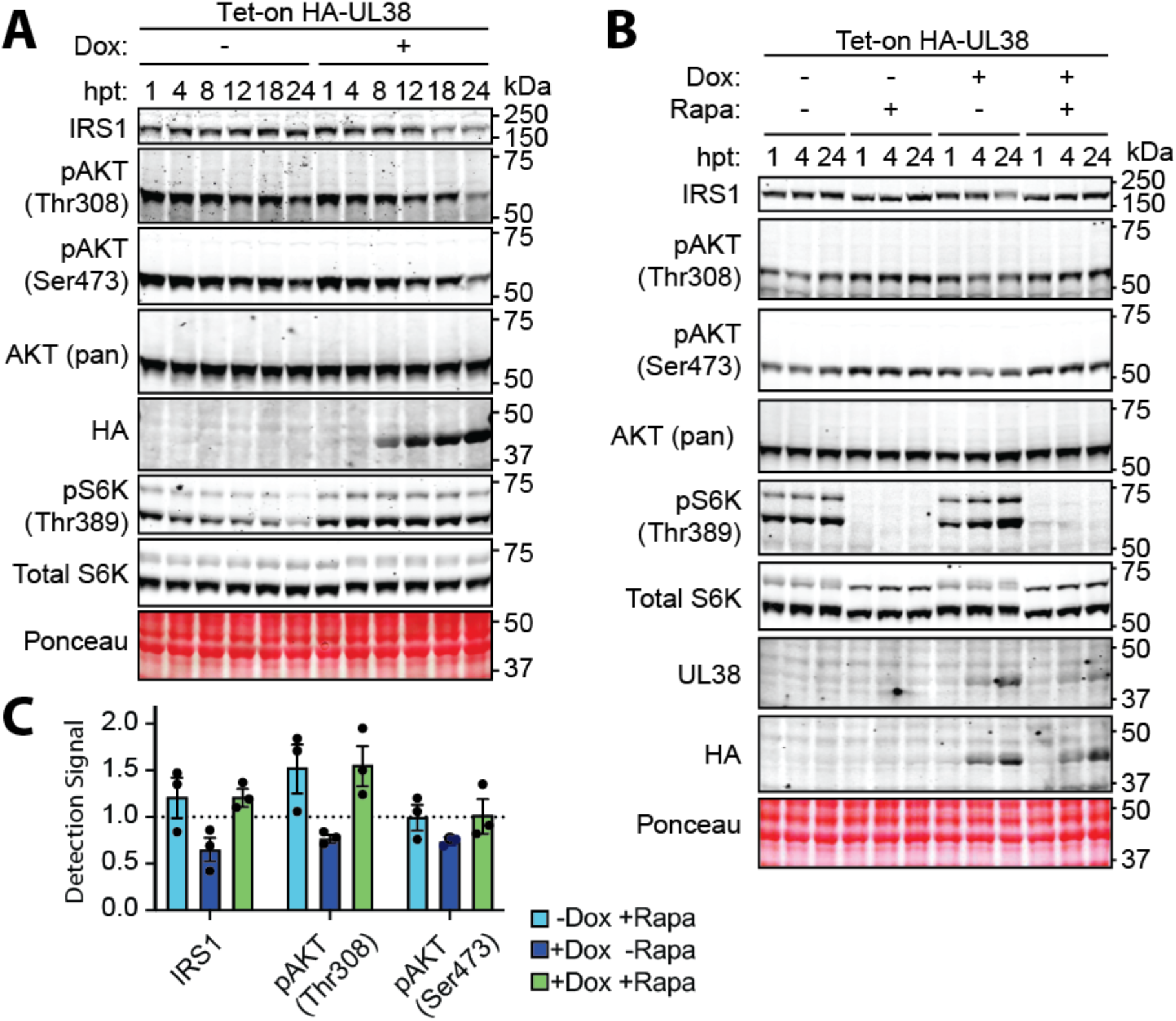
UL38 is sufficient to inactivate AKT. (**A-C**) Fibroblasts transduced with a lentivirus harboring a “Tet-on” HA-UL38 expression cassette were either induced for UL38 expression using 100 ng/mL doxycycline (Dox, +) or carrier-alone (-) in the presence of serum. Lysates were taken at indicated times post-treatment (h post treatment, hpt) and analyzed by Western blotting for the indicated proteins and phospho-epitopes. Ponceau stain was used to monitor protein loading and to ensure efficient transfer to nitrocellulose membranes. (**B-C**) Cells were either Dox induced or mock-induced using carrier alone, and treated or not with 50 nM rapamycin (Rapa) prior to lysis, as indicated. (**C**) At 24 hpt, the signal intensity from far-red flurophore conjugated antibodies detecting phospho-specific bands or total IRS1 protein were quantified and normalized to those from detection of AKT (pan). Results were graphed relative to a baseline signal of 1.0 determined for untreated control cells (dotted line). The arithmetic mean was calculated and graphed. Tukey’s multiple comparisons statistical test was used to compare each condition to each other and labeled with asterisks (ns=P>0.05, *=P<0.05, **=P<0.01, ***=P<0.001, ****P<0.0001). Error bars=SEM. n=3.

If UL38 was sufficient to cause mTORC1-dependent IRS1 degradation and subsequently, AKT inactivation, then the IRS1 destabilization and loss of AKT responsiveness to serum we observed in cells expressing UL38 should be reversed upon rapamycin treatment. To test this, we induced UL38 expression in the presence of serum, and then treated with rapamycin (or carrier-alone) over a time course from 1 h to 24 h following Dox induction. After inducing HA-UL38 expression in fibroblasts that had been stably transduced with a “Tet-on” HA-UL38 cassette, we detected UL38 protein accumulation, both by a UL38-specific monoclonal antibody, and by an antibody specific for the HA tag (**FIG 7B**). Concomitantly, we detected increases in pS6K (Thr389), as expected due to UL38-mediated activation of mTORC1 (**FIG 7B**). We also observed a 35.5% reduction in IRS1 detection by 24 h post Dox induction. However, rapamycin treatment rescued IRS1 abundance, showing over 2-fold (2.2×) higher levels relative to the Dox-induced comparator lacking rapamycin treatment We also observed that the IRS1 species in the Dox-induced (UL38 expression) condition showed decreased mobility in SDS-PAGEd compared to uninduced +Rapa and Dox induced/+Rapa controls. Furthermore, in the Dox-induced condition, we saw a 32% decrease in the detection signal for AKT Thr308 phospho-epitope and a 36% decrease in that for AKT phospho-Ser473. However, treatment with rapamycin rescued the detection of these phospho-epitopes, respectively, by up to 156% and 103% relative to the levels in non-inhibited controls (**FIG 7B–C**). These data indicate that UL38 expression is sufficient to trigger mTORC1-dependent IRS1 degradation and downstream inactivation of AKT.

Finally, we tested whether UL38 is necessary to destabilize IRS1 and render AKT non-responsive to serum in the authentic context of HCMV infection. We therefore infected cells in the presence of serum using a UL38-null mutant of HCMV strain AD169 (AD ΔUL38) <u>[72]</u> alongside parental wild-type (WT) AD169 and mock infection controls, and gathered lysates over a 24 h infection time course. We focused on this time interval because our data show that within 9 hpi to 12 hpi of HCMV infection, AKT accumulates in a form that is hypophosphorylated at its two key activating residues (Ser473, Thr308) and insensitive to serum stimulation (**FIG 6B**), <u>[42]</u>.

As expected, expression of UL38 was observed in cells infected with WT HCMV strain AD169 but not in cells infected with a UL38-null mutant strain AD169 (ΔUL38) comparator virus. Moreover, the kinetics of UL38 expression in WT HCMV infected cells tightly correlated with increased detection of pS6K (Thr389) (**FIG 8A**). This observation is consistent with the established roles of UL38 as a viral activator of mTORC1 <u>[46,48]</u>. Levels of the HCMV proteins UL44 and IE1 were indistinguishable between WT and ΔUL38 virus infections, suggesting that overall progression through the viral lytic cycle was comparable across the two settings. (**FIG 8A**). However, we did detect a marked decrease in IRS1 protein abundance in WT virus-infected cells that was not observed in ΔUL38 mutant virus or mock infection settings. We quantified 24 hpi IRS1 detection signals from three independent biological replicates and were able to estimate that WT HCMV-infected cells undergo a 50% decrease in IRS1 abundance relative to mock-infected cells (FIG 8B). In contrast, UL38-null infected cells, on average, showed a 22% increase in IRS1 levels relative to mock-infected cells. IRS1 from WT-infected cells, but not mock or ΔUL38-infected cells, also showed decreased mobility. This effect was most striking at 12 hpi and 24 hpi in SDS-PAGE. The IRS1 mobility differences seen during WT infection were accompanied by 86% and 82% reductions in the detection signals for phosphorylation at AKT Thr308 and Ser473, respectively, compared to mock infection. In ΔUL38-infected cells, these decreases in AKT phosphorylation (relative to mock) were found to be only 47% and 58% respectively (FIG 8A–B). We interpret from these results that UL38 is required for IRS1 degradation and inactivation of AKT during infection. Taken together, our findings identify UL38 as the major viral gene product that causes IRS1 degradation and renders AKT insensitive to serum growth factors during HCMV infection.

**Figure 8:**
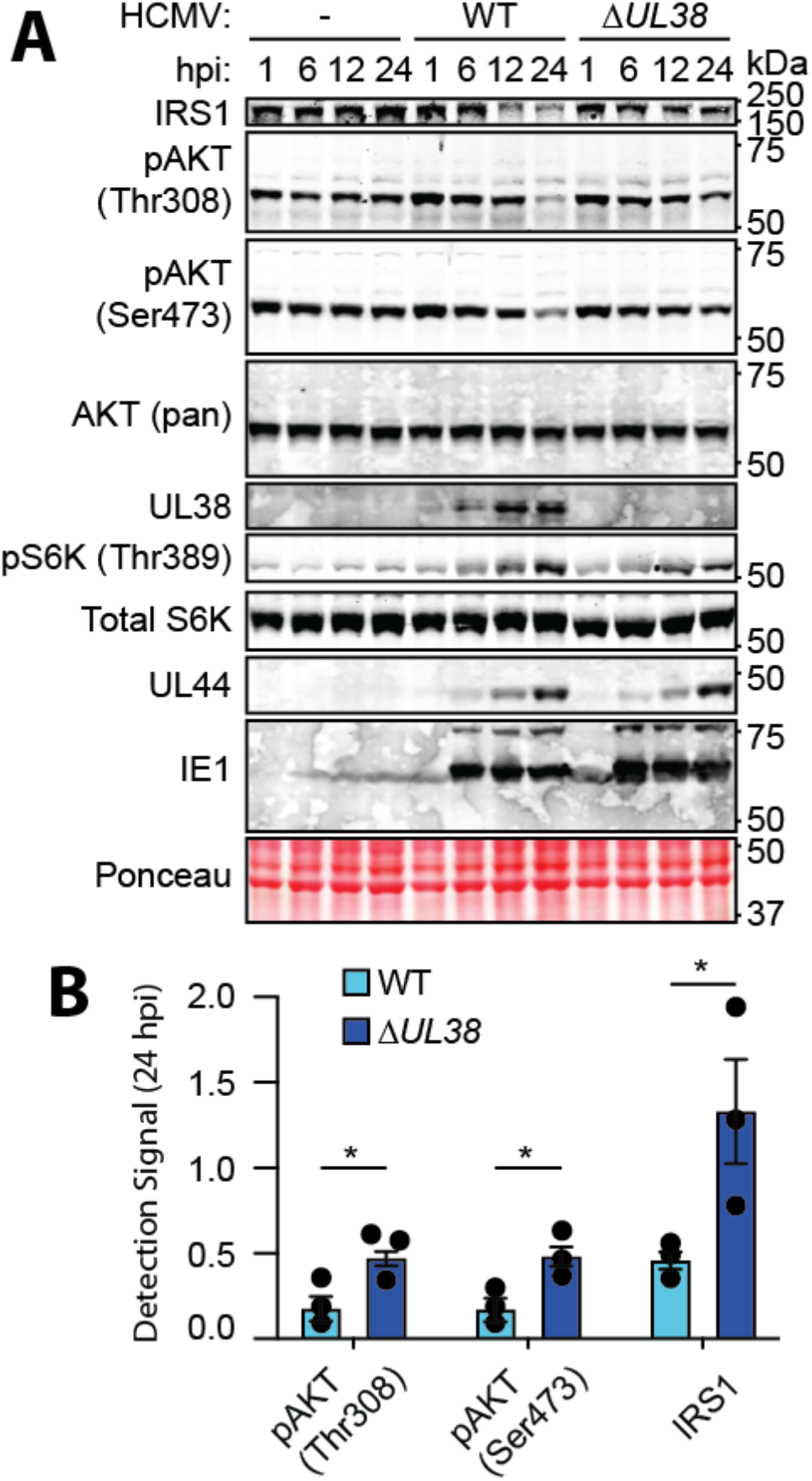
UL38 inactivates AKT during HCMV infection. (**A-B**) Fibroblasts were mock-infected or infected with wild-type (WT) HCMV strain AD169 or an AD169 whose UL38 gene locus has been genetically knocked out for **UL38** (ΔUL38) (MOI=2 TCID50/cell). At 1, 6, 12, and 24 h post-infection (hpi), lysates were gathered and a Western blot analysis was carried out to detect the indicated proteins and phospho-epitopes. Ponceau S stain was used to indicate overall protein loading and to ensure efficient transfer to nitrocellulose membranes. (**B**) Signals from detection of the indicated phospho-specific bands and IRS1 were quantified for 24 hpi samples and normalized to signal from detection of AKT (pan). The arithmetic mean was calculated and graphed. Sidak statistical test were used to compare each condition to the control and labeled with asterisks (ns=P>0.05, *=P<0.05, **=P<0.01, ***=P<0.001, ****P<0.0001). Error bars=SEM. n=3.

## DISCUSSION

How intracellular pathogens commandeer the metabolic resources and nutrients from host cells is central to their biology and strategies for persistence. The PI3K/AKT/mTOR pathway is a key battleground in the tug of war to usurp the host cell’s macromolecular synthesis machinery. Unlike many other viruses, HCMV inactivates AKT during infection, and how this occurs has remained unknown. Our study reveals that HCMV relies on a canonical, cell-intrinsic negative feedback loop to render AKT insensitive to serum. Given that UL38 is a powerful activator of mTORC1 [46–48], this mechanism should not be surprising [73,74]. Nonetheless, there has been much confusion and controversy concerning how HCMV rewires host cell signaling, particularly regarding AKT and mTOR [75,76]. Indeed, HCMV infected cells have a penchant for defying expected results. For example, the mTORC1 inhibitor rapamycin ordinarily prevents phosphorylation of both S6K and 4E-BP1. However, during HCMV infection 4E-BP1 phosphorylation occurs independently of mTORC1 [41].

Like many other intracellular pathogens, herpesviruses require cap-dependent mRNA translation to replicate, and therefore deploy mechanisms to maintain mTORC1 activity [77]. In fact, many herpesviruses directly activate PI3K/AKT, relying on virally encoded proteins to do so, e.g. K1 of KSHV [48,78,79], ORF12 of VZV [80], and LMP2A/LMP1 of EBV [81–83]. The herpes simplex virus 1 (HSV-1) tegument protein VP11/12 activates AKT [84,85], while viral protein kinase US3 mimics AKT [1]. In contrast, during the HCMV lytic cycle, the virus rapidly renders AKT inactive and insensitive to serum (**FIG 6B**) [43]. Because the HCMV immediate-early protein UL38 directly sustains mTORC1 activity [48], cap-dependent translation is maintained independently of AKT. Moreover, expression of constitutively active AKT is detrimental to lytic cycle progression [42], apparently because this prevents nuclear localization of FoxO transcription factors (e.g., FoxO3a) that play key roles in viral replication [86,87].

Although we have focused here on deciphering how HCMV inactivates AKT during lytic infection, our results may have implications for HCMV latency. Pharmacological inhibition of PI3K/AKT is among the most broadly reliable methods to reactivate HCMV from latent carriage [43] [reviewed in [88]]. Moreover, HCMV proteins, such as UL138 and UL135, that regulate EGFR–a key upstream modulator of PI3K/AKT, play pivotal yet opposing roles in maintenance of latency [44,89]. Because HCMV must render AKT inactive to efficiently replicate [42], UL38 likely plays a major role in the latent-lytic switch. Intriguingly, translation of viral messages is initially disfavored over host transcripts [90], yet UL38 profoundly impacts the translational milieu to favor translation of viral messages [91]. Given that latently-infected cells show surprisingly broad, largely unrestricted transcription of viral mRNAs, albeit at low levels [92,93], the mechanisms that govern the accumulation of UL38 protein during establishment and reactivation from latency merit further study.

## MATERIALS AND METHODS

### Recombinant DNA

All novel plasmids were generated using Gibson Assembly of PCR-amplified sequences into digested backbone plasmids. Before using newly constructed plasmids in experiments, they were sequence-confirmed by long read Oxford Nanopore long read sequencing (SNPsaurus, Eugene, OR) and/or Sanger sequencing (Genewiz) [data not shown]. Unless otherwise noted, all oligonucleotide primers were synthesized by Integrated DNA Technologies (IDT). To construct a lentivirus vector for constitutive expression of HA tagged UL38, pLV HA-UL38-IRES-Hygro, an HA-epitope tagged UL38 ORF was amplified from PB-TAC-ERP2_HA_UL38 using Gib.BamHI.pLVhT.UL38_Fw (CTTCCATTTCAGGTGTCGTGAGACCGGTGCCACCATGTACCCATACGA) and Gib.EcoRI.pLVhT.UL38_Rv (TAGAGCGGCCGCCCTCGAGGACGCGTCTAGACCACGACCACCATCTG) primers. The polymerase chain reaction (PCR) was run using 0.2 U/μL of KOD Hot Start DNA polymerase (Novagen, Cat# 71316), 0.3 μM each of primers, 1.3 M betaine monohydrate [final] (Sigma Prod. #: B2754), 0.2 μM dNTPs (Novagen, Cat#: 71154), 1X KOD Hot Start DNA Polymerase Buffer (Novagen, Cat#: 71155), 1.5 μM MgSO_4_ (Novagen Cat#: 71156), 1 ng of template and water to 50μL. The PCR product was run out on a 1% agarose (GoldBio Cat#: A-201-1000) gel in Tris-Acetate-EDTA (TE) pH 8.5 (40 mM Tris base, 20 mM glacial acetic acid, 1 mM EDTA). A pLV IRES-Hygro plasmid (Addgene #85140) was digested with BamHI (NEB Cat#: R3136S) and EcoRI (NEB Cat#: R3101S) in 1x CutSmart buffer (NEB Cat#: B7204S) to release the insert, and the 9.126 kb backbone was gel purified using a Qiaex II Gel Extraction Kit (Qiagen Cat. No. 20051), alongside the PCR product, as per manufacturer instructions. The PCR product and pLV IRES-Hygro plasmid backbone were assembled using HiFi DNA Assembly Master Mix (NEB Cat#: M5520AA) according to the manufacturer’s protocol. NEB Stable Competent E. coli (NEB Cat#: C3040H) were transformed and plated on LB Agar (Fisher BioReagents Cat#: BP1425-500) containing 50 µg/mL carbenicillin (GoldBio Cat#: C-103-5) and incubated overnight at 37°C. The next day, colonies were picked and grown in LB broth (Fisher Cat#: BP1426-2) containing carbenicillin. To construct a “Tet on” *UL38* lentivirus vector, the HA-*UL38* sequence from the pLV-HA-UL38-IRES-Hygro was amplified using pOUPc_HA_UL38_FWD (CCACTTCCTACCCTCGTAAACCGGTGCCACCATGTAC) and pOUPc_UL38_REV (GGAGGCCAGATCTTAACGCGTCTAGACCACGACCACCATCTG) primers. A pOUPc plasmid (e.g., pOUPc-UL148^HA^, ref: [94] was linearized by double-digested with MluI-HF (NEB Cat#: R3198L) and AgeI-HF (NEB Cat#: R3552L), and the 11.787 kb backbone fragment was isolated by agarose gel electrophoresis, gel purified, and Gibson assembled with the HA-UL38 PCR product.

### Cells and Viruses

Primary human foreskin fibroblasts (HFFs) (ATCC SCRC-1041) were transduced with a lentivirus encoding human telomerase reverse transcriptase (hTERT) to generate telomerase-immortalized HFF cells (HFFT). The adult retinal pigment epithelial cell line ARPE-19 was purchased from ATCC (ATCC CRL-2302). HFFT and ARPE-19 cells were grown in Dulbecco’s modified Eagle’s medium (DMEM) (Genesee Scientific, Cat # 25-500) supplemented with 25 μg/mL of gentamicin (Gibco cat # 15750-060), 10 μg/mL of ciprofloxacin HCl (GenHunter SKU: Q901-1G) and 5% Newborn Calf Serum (GeminiBio, Cat #. 100-504). HeLa cells (American Type Culture Collection [ATCC], Cat # CCL-2) and human embryonic kidney (HEK) 293T cells (a kind gift from Victor DeFillippis, Oregon Health Sciences University) were grown in Opti-MEM GlutaMAX media (Gibco Cat # 51985-034) supplemented with ciprofloxacin and gentamicin (as above) and 10% Fetal Bovine Serum (GeminiBio, Cat # 900-208). All cells were incubated at 37°C in a humidified 5% CO_2_ incubator for routine growth and for HCMV infection studies.

Stocks of HCMV strains Towne (ATCC VR-977) and parental AD169rv [95] were produced by low MOI infection of HFFTs (MOI of 0.01 TCID_50_/cell). Strain AD169rv repaired for expression of UL131 (AD169r131) [96] was amplified in ARPE-19 epithelial cells by infecting at an MOI of 0.01 TCID_50_/cell. Approximately 4-6 days after complete cytopathic effect (CPE) was observed, cell culture media and cells were collected and the cells were homogenized using 10 strokes of a Dounce homogenizer (Wheaton Cat # 357542). The media/homogenate was cleared of cell debris by centrifugation at 2000 rpm for 10 min. A sorbitol (20% D-sorbitol, 25 mM HEPES pH 7.5, 1 mM MgCl_2_, 100 μg/mL Bacitracin) underlay was gently pipetted under the supernatant in polyallomer ultracentrifuge tubes (Beckman-Coulter, Cat #: 326823). The virus was ultracentrifuge concentrated using a SW32 Ti swinging bucket rotor 24,000 rpm for 2 h at 4°C) in a Beckman Coulter Optima L-90K Ultracentrifuge. The supernatant was removed, and virion pellets were resuspended in 100% bovine serum albumin fraction V (REF: 15260-037) and stored at –80°C until use.

To produce stocks of ADΔUL38, a *UL38*-null HCMV strain AD169, HFFTs constitutively expressing a HA-UL38-IRES-Hygro (see below) were infected with AD169 dlUL38 (a generous gift from Dr. Joshua Munger, University of Rochester) at MOI 0.01 TCID50/cell <u>[72]</u>. Virus was harvested once infected cell monolayers showed complete CPE and was concentrated, resuspended and stored as described above.

### Virus Titration

The infectious titers of viral stocks were measured on HFFT by end point dilution to determine the 50% tissue culture infectious dose (TCID_50_), as previously described [97]. Briefly, 1:10 serial dilutions were prepared in 5% NCS DMEM with antibiotics, and 8 wells were infected per each dilution in a 96-well cluster plate format. 10 d post-infection, wells were scored positive or negative for viral plaque formation/cytopathic effects and the viral titer was calculated using the Spearman-Karber method [98,99]

### Serum and insulin response studies

For stimulation of HFFTs with insulin and/or serum, cells were serum-starved overnight with serum-free DMEM supplemented with 25 μg/mL of gentamicin and 10 μg/mL of ciprofloxacin HCl. Media was replaced with DMEM containing gentamicin and ciprofloxacin HCl alone or containing antibiotics and 10% NCS and/or 200 ng/mL Recombinant Human Insulin growth factor 1 (R&D Systems Cat#: 291-G1) for indicated minpost-treatment. Cells were then washed with PBS and lysed as described below for Western blot analysis.

For serum responsiveness assays, HFFTs were serum-starved overnight with serum-free DMEM, supplemented only with antibiotics. Cells were then infected with HCMV strain Towne (MOI=2 TCID_50_/cell) or mock-infected in the same serum-free media the next morning for the indicated times. 10 min prior to washing and lysis (see below), cells were serum stimulated by gently adding 10% NCS [final] directly to the media.

### Western blotting

Cells were washed once with Dulbecco’s phosphate buffered saline (PBS) (Genesee Scientific REF: 25-508), lysed in 1x Laemmli Buffer containing 1x Protease Inhibitor Cocktail (APExBio, Cat. # K1007) and 1x Phosphatase Inhibitor Cocktail 1 (APExBio, Cat. # K1012), collected and stored at –20°C until analysis. 2-Mercaptoethanol (Sigma-Aldrich, Cat. # M7154) was added for the final 5% vol/vol and lysates were incubatedat 65°C for 10 mins. Lysates were then vortexed for 1 min at max speed and loaded into a 10% sodium dodecyl sulfate (SDS)-polyacrylamide gel and run at constant 100 V for 1 h 45 min in running buffer (25 mM Tris, 190 mM Glycine, 3.46 mM SDS) alongside a Precision Plus Protein Dual Color Standard (Bio-Rad, Cat #1610374) in. Proteins were transferred to a 0.45 μm nitrocellulose membrane (Amersham Protran, Cytiva Cat #10600007) constant 20V for overnight in transfer buffer (25 mM Tris, 190 mM glycine, 20% Methanol). Membranes were soaked in 0.2% Ponceau S Stain ((Sigma-Aldrich, Cat. # P3504) weight/vol, 5% vol/vol glacial acetic acid (J.T. Baker 9508-06)) and imaged on ESPON Perfection V700 Photo using EPSON Scan (Ver 5.1.0f0). Membranes were washed in 5% bovine serum albumin, fraction V (BSA) (Fisher Scientific BP1605) w/v diluted in PBS (137 mM NaCl, 2.7mM KCl, 10mM Na_2_HPO_4_, 1.8 mM KH_2_PO_4_) for 1 hour. Primary and secondary antibodies were added at indicated dilution (see antibodies section) for 10 min and each antibody was washed four times with PBS containing 0.5% TWEEN-20 (Fisher Scientific, Cat. # BP337) v/v (PBST) for 1 hour total. The membrane was scanned on a LI-COR Odyssey CLx Infrared Imaging System. Quantification was performed using Image Studio (Ver 5.2). Images were cropped, straightened, and uniformly processed using Adobe Photoshop CS5 (Ver 12.0) and labeled using Adobe Illustrator CS5 (Ver 399) for Macintosh.

### Membrane fractionation studies

HFFTs were serum starved overnight and then infected with Towne (MOI=2 TCID_50_/cell, or mock infected. At 12 h post-infection (hpi), cells were either subjected to continued serum starvation or treated with 5% NCS for 30 min. Cells were then washed with ice-cold PBS and scraped into Fractionation Buffer (25 mM Tris-HCl (pH 7.4), 2 mM EDTA, 10 mM NaCl, and 0.25 M sucrose) containing 1x Protease Inhibitor and 1x Phosphatase Inhibitor Cocktail I (ApexBio). Cells were allowed to swell for 10 min on ice and then homogenized with 30 strokes of a glass Dounce homogenizer. An aliquot of lysate was then taken and reserved for use as “Total Lysate,” and the remainder was centrifuged at 2,000 x g for 5 min at 4°C. The supernatant was decanted into 0.8 mL Ultra-Clear Tubes (Beckman Coulter, Cat# 344090) and inserted into a split adapter (Beckman Coulter, Cat# 356860) within swinging buckets of an SW55Ti rotor (Beckman Coulter, Cat# 342194). After centrifugation at 104,000 x g for 30 min at 4°C using a Beckman Coulter Optima L-90K Ultracentrifuge, the supernatant was collected and treated as the “Cytoplasmic Fraction” and the remaining pellet was resuspended in Fractionation Buffer and Labeled “Membrane Fraction”. For all samples, 2-Mercaptoethanol was then added to 5% (vol/vol) and Laemmli Buffer was added to 1x, and processed for SDS-PAGE and Western blot analysis as described above.

### Antibodies

The antibodies used for Western blots in this study are shown in **Table 1**.

**TABLE 1.**

|  | Antibody Name | Species | Source | Cat# | Dilution |
| --- | --- | --- | --- | --- | --- |
| **Primary** | AKT (pan) (40D4) | Mouse | Cell Signaling Tech | 2920S | 1:2000 |
|  | PRAS40 (D23C7) XP | Rabbit | Cell Signaling Tech | 2691S | 1:1000 |
|  | Phospho-PRAS40 (Thr246) (D4D2) XP | Rabbit | Cell Signaling Tech | 13175S | 1:1000 |
|  | p70 S6 Kinase (49D7) | Rabbit | Cell Signaling Tech | 2708S | 1:1000 |
|  | Phospho-p70 S6 Kinase (Thr389) (108D2) | Rabbit | Cell Signaling Tech | 9234S | 1:1000 |
|  | Phospho-AKT (Ser473) (D9E) XP | Rabbit | Cell Signaling Tech | 4060S | 1:1000 |
|  | Phospho-AKT (Thr308) (D25E6) XP | Rabbit | Cell Signaling Tech | 13038S | 1:1000 |
|  | P-S6 Ribosomal Protein (Ser235/236) (D57.2.2E) XP | Rabbit | Cell Signaling Tech | 4858S | 1:1000 |
|  | S6 Ribosomal Protein (5G10) | Rabbit | Cell Signaling Tech | 2217S | 1:1000 |
|  | p-PDK1 (S241) (C49H2) | Rabbit | Cell Signaling Tech | 3438T | 1:1000 |
|  | Phospho-PTEN (Ser380/Thr382/383) | Rabbit | Cell Signaling Tech | 9554S | 1:1000 |
|  | PTEN (D3Q6G) Mouse mAb #14642 | Mouse | Cell Signaling Tech | 14642S | 1:1000 |
|  | IRS-1 Antibody #2382 | Rabbit | Cell Signaling Tech | 2382S | 1:1000 |
|  | 62-X-IE1 | Mouse | Tom Shenk | n/a | 1:1000 |
|  | 308 anti-UL38 | Mouse | John Purdy | n/a | 1:200 |
|  | Integrin B1 (TS2/16) | Mouse | Santa Cruz | sc-53711 | 1:200 |
|  | HA | Mouse | Santa Cruz | sc-7392 | 1:200 |
|  | GAPDH | Mouse | Proteintech | 60004-1-Ig | 1:20000 |
|  | UL44 clone CH13 | Mouse | Virusys | P1202-1 | 1:500 |
| **Secondary** | IRDye 800CW | Donkey anti-rabbit | LI-COR | 926-32213 | 1:12000 |
|  | IRDye 680RD | Goat anti-mouse | LI-COR | 926-68070 | 1:12000 |

### HeLa cell infection

For Western blot analysis of AKT activity of HCMV infected HeLa cells, HeLa cells were seeded in 10% FBS OptiMEM supplemented with 25 μg/mL of gentamicin and 10 μg/mL of ciprofloxacin HCl overnight. Cells were infected with HCMV strain AD169r131 (MOI=2 TCID_50_/cell) and a Western blot analysis was carried out as described above.

### Transfection

Transfection grade plasmid DNAs for this study were prepared using Nucleobond Xtra Midi kits, according to the manufacturer’s instructions (Machery-Nagel, Inc., Allentown, PA). HEK 293Ts were reverse-transfected with pcDNA3-AKT-PH-GFP (Addgene #18836) or pcDNA3-AKT-PH[R25C]-GFP (Addgene #18837), (a gift from Dr. Tamas Balla, National Institutes of Child Health and Human Development) [57]. 3 μg of plasmid DNA and 1μg/mL of linear transfection-grade PEI in sterile PBS were mixed with HEK-293T cells, which were then allowed to attach to 35mm glass bottom culture dishes (MatTek Part Number: P35G-1.5-14-C) in 10% FBS OptiMEM overnight at 37°C, 5% CO_2_ in a humidified incubator. Cells were then mock-infected or infected with HCMV strain AD169r131 (MOI=2 TCID_50_/cell), as described above.

### Live Cell Imaging

Two hours prior to imaging, cell culture media was replaced with serum-starvation media (DMEM containing no serum). Cells were mock-treated or treated with 10% FBS (final) and imaged at 10 minutes post-treatment. During serum stimulation and imaging, the cells were housed in a 37°C humidified 5% CO_2_ stage-top incubation chamber (Tokai Hit Chamber: WSKMX & controller: STXF, Tokai Hit Co, Ltd, Shizuoka-ken, Japan). Images were obtained on an Olympus CSU W1 Spinning Disk Confocal microscope (Olympus Life Science) with a sCMOS camera (Hamamatsu Fusion) using a 60x oil immersion objective (Olympus UPlanApo 60x).

### Membrane recruitment studies

Images were imported into ImageJ 1.45s (Java 1.6.0_65) and for each cell, 50 points along the outer membrane and 50 points within the cell outside of the nucleus were selected and measured for pixel intensity. The average signal measured along the membrane was normalized to the average signal within the cell. Five independent biological replicates were carried out and the standard error of the mean (SEM) was propagated. The normalized average pixel density at the membrane was graphed using GraphPad Prism for MacOS, Ver 9.5.0 (DotMatics, Inc.) with error bars representing SEM. Tukey’s multiple comparisons statistical test was used to compare each condition to each other and labeled with asterisks (ns=P>0.05, *=P<0.05, **=P<0.01, ***=P<0.001, ****P<0.0001).

### UV-inactivation of virions

For FIG 5, HCMV strain Towne (MOI=2 TCID_50_/cell) was placed on ice for 5 minutes and either mock-treated or treated with 125 mJ of UV light for 0, 15, 30, 45 or 60 seconds (Bio-Rad GS Gene Linker UV Chamber). The inoculum was placed on fibroblasts for 12 hours, lysates were gathered and a Western blot analysis was run as described above. For figure 4 B-D, HCMV strain Towne (MOI=2 TCID_50_/cell) was placed on ice for 5 minutes and either mock-treated or treated with 125 mJ of UV light for 60 seconds. Lysates were gathered at indicated time points and a Western blot analysis was run as described above.

### Lentiviral vector transduction

Human embryonic kidney 293 cells transformed with T-antigen (HEK-293T) were reverse-transfected with the pOUPc-HA-UL38, or pLV HA-UL38-IRES-Hygro plasmid using linear polyethyleneimine (PEI) (Polysciences, Cat.# 23966). Briefly, subconfluent HEK-293T cells were reverse-transfected with 56.7 μg of transfer plasmid, 21.3 μg of psPAX2, 7.1 μg pMD2.G, and 1 μg/mL of PEI in sterile PBS. Cells were mixed with the transfection mixture and seeded on two 15 cm dishes in 10% FBS OptiMEM overnight. The next morning, the media was replaced with OptiMEM containing 10% FBS. After 2 d, the media supernatant was collected, filtered through a 0.45-micron syringe filter (Corning Cat # 431220) and used to transduce low-passage HFFTs supplemented with 8 μg/mL of polybrene. 48 h post-transduction, cells were subjected to selection with either 1 μg/mL of puromycin or 50 μg/mL of hygromycin, according to the selectable marker expressed by the lentiviral vector (GoldBio, Cat#: H-270-1). Dox-inducible myrAKT-HA fibroblasts (**FIG 4**) were generated as described previously [42].

### Drug Treatments

For the experiment shown in Figure 4, HFFTs stably transduced with a previously described “tet-on” myrAKT-HA expression cassette [42] were seeded overnight in media containing 5% NCS and 100 ng/mL of doxycycline (Dox). The cells were then infected with HCMV strain Towne (MOI=2 TCID_50_/cell) in 5% NCS media containing Dox (100 ng/mL) and incubated for the indicated times (37°C, 5% CO_2_) until harvest for for Western blot analysis (as described above).

As with Figure 4, for the experiment shown in Figure 7, HFFT cells stably transduced with a constitutive UL38-HA expression cassette were seeded overnight in media containing 5% NCS. The next morning, cells were then treated with 100 ng/mL of Dox. For the experiment shown in Figure 7B, 50 nM rapamycin (rapa) was added at the same time of Dox addition. The cells were exposed to the drug(s) until harvest for Western blot analysis, as described above.

For the experiment shown in Figure 6D, HFFT were serum starved overnight in DMEM lacking serum, and then infected with HCMV strain Towne (MOI=2 TCID_50_/cell). At 3 hpi, cells were treated with 50 nM rapa in DMEM lacking serum. At 24 hpi, serum (10% NCS, final) was added (in the presence of50 nM rapa DMEM) for a 10 min incubation at 37°C, 5% CO_2_, after which lysates were collected for Western blot analysis, as described above.

### Western blot quantification and data analysis

All graphs were generated in GraphPad Prism by DotMatics for MacOS (Ver 9.5.1) and analyzed using built-in statistical analysis tools. Sidak and Tukey’s multiple comparisons statistical tests were used, as indicated in Figure legends, to compare each condition to each other and labeled with asterisks (ns=P>0.05, *=P<0.05, **=P<0.01, ***=P<0.001, ****P<0.0001). Far-red fluorescent dye signals from secondary antibodies on Western blots were acquired on a Li-Cor Odyssey CLx, and the smallest possible rectangular box that contained the band of interest was drawn and repeatedly copied to the neighboring bands to maintain size consistency. Detection signal values for each band were exportedd to a spreadsheet, and phospho-specific antibody band values (e.g., phospho-AKT Ser473) were normalized to cognate protein detection signals (e.g., pan AKT), or in the case of IRS1, to AKT detection signal (as a loading control).

For serum responsiveness studies, values were then normalized to “+Serum”/ “NoVirus” controls. The average (arithmetic mean) of results from three independent biological replicates was calculated, graphed, and analyzed in GraphPad Prism 9.5.1. For membrane fractionation studies, total AKT quantification values in the membrane fraction were normalized to total lysate values then all conditions were subsequently normalized to “No Serum” controls. The arithmetic means of values from four independent biological replicate experiments were calculated, statistical analysis was performed, and final values were graphed. For experiments on UV-inactivated virions, detection signals for phospho-epitopes relative to detection signal for the cognate protein at 24 hpi for both conditions were normalized to phospho-epitope signals relative to cognate protein signal at 1 hpi. The arithmetic mean of results from three independent biological replicates was graphed. For time course studies, the normalized detection values for phosphospecific bands were further normalized to mock-infected controls at 1 hpi. For rapamycin experiments on infected cells, normalized detection values from infected lysates were normalized to mock infection/ “No Rapamycin” controls, and further normalized to HCMV infection/ “No Rapamycin” controls. Averages (arithmetic mean) of three independent biological replicates were calculated, statistical analysis was performed, and values were graphed. For rapamycin inhibition experiments on myr-AKT “tet-on” inducible cells, normalized detection values at 24 h post induction were normalized to “No Doxycycline”/ “No Rapamycin” controls. Average (arithmetic mean) values from three independent biological replicates were calculated and graphed. For experiments involving UL38-null HCMV (ADΔUL38), values normalized to AKT loading control at 24 hpi were further normalized to 1 hpi. Averages (arithmetic mean) of three independent biological replicates were calculated and graphed.

## ACKNOWLEDGEMENTS.

We thank John G. Purdy (University of Arizona) and Thomas E. Shenk (Princeton University) for providing UL38 monoclonal antibodies, and Matthew Raymonda, Xenia Schafer and Joshua C. Munger of the University of Rochester (Rochester, New York) and Thomas E. Shenk (Princeton University) for sharing with us the ADΔUL38 virus. We also thank Malgorzata Bienkowska-Haba of the LSU Health Sciences Center Shreveport Microscopy Core for her expert assistance with live cell imaging. We are grateful to Stanimir S. Ivanov (LSU Health Shreveport) for helpful discussions. Research reported in this publication was supported by the National Institute of Allergy and Infectious Diseases and the National Institute of General Medical Sciences of the National Institutes of Health under award numbers R01-AI143191 (to F.D.G., N.J.M., and J.P.K.) and P20-GM134974. The content is solely the responsibility of the authors and does not necessarily represent the official views of the National Institutes of Health.

